# *Xenopus* Ssbp2 is required for embryonic pronephros morphogenesis and terminal differentiation

**DOI:** 10.1101/2023.04.15.537039

**Authors:** Ailen S. Cervino, Mariano G. Collodel, Ivan A. Lopez, Daniel Hochbaum, Neil A. Hukriede, M. Cecilia Cirio

## Abstract

The nephron, functional unit of the vertebrate kidney, is specialized in metabolic wastes excretion and body fluids osmoregulation. Given the high evolutionary conservation of gene expression and segmentation patterning between mammalian and amphibian nephrons, the *Xenopus laevis* pronephric kidney offers a simplified model for studying nephrogenesis. The Lhx1 transcription factor plays several roles during embryogenesis, regulating target genes expression by forming multiprotein complexes with LIM binding protein 1 (Ldb1). However, few Lhx1-Ldb1 cofactors have been identified for kidney organogenesis. By tandem-affinity purification from kidney-induced *Xenopus* animal caps, we identified <u>s</u>ingle-<u>s</u>tranded DNA <u>b</u>inding <u>p</u>rotein <u>2</u> (Ssbp2) interacts with the Ldb1-Lhx1 complex. Ssbp2 is expressed in the *Xenopus* pronephros, and knockdown prevents normal morphogenesis and differentiation of the glomus and the convoluted renal tubules. We demonstrate a role for a member of the Ssbp family in kidney organogenesis and provide evidence of a fundamental function for the Ldb1-Lhx1-Ssbp transcriptional complexes in embryonic development.

## Introduction

The vertebrate kidney is a specialized excretory organ with a key role in osmoregulation and metabolic and drugs waste removal. During development, the kidney emerges bilaterally from the intermediate mesoderm and develops through a series of progressively more complex forms, the pronephros, mesonephros and metanephros, the later only found in mammals, birds, and reptiles. All of them share the same basic structural and functional unit called the nephron consisting of a filtration unit, the glomerulus in integrated and the glomus in non-integrated nephrons, the renal tubule, subdivided into proximal and distal segments and the connecting tubule ^1, 2^. Formation of each kidney is induced by the preceding nephric tissue leading to an increased organizational complexity ^3, 4^. While the functional pronephros of amphibian and fish embryos is composed of two nephrons, the metanephros of the adult amniotes is composed of many nephrons, approximately 1 million in humans ^5^. Despite anatomical differences, many genes and processes that govern pronephros formation are conserved in individual metanephric nephrons development ^6–8^. In fact, the functional *Xenopus laevis* pronephros offers a simplified model for the study of nephron development and human kidney diseases ^9–12^.

Signaling pathways and transcriptional regulators involved in early kidney specification have been systematically investigated, however less is known about downstream effectors required for nephric patterning and morphogenesis. In this regard, the role of the LIM-class homeobox transcription factor Lhx1 is essential for kidney development ^13–16^. However, in this tissue, few Lhx1 interacting proteins and downstream targets have been identified ^16, 17^. Lhx1 binding to <u>L</u>IM <u>d</u>omain <u>b</u>inding protein 1, Ldb1, through the LIM domains, triggers activation and formation of a tetrameric base-complex. Together, they regulate gene expression through the formation of multiprotein transcriptional complexes with crucial roles during development ^14–16, 18–20^. Previously, we identified Ldb1-Lhx1 binding proteins by tandem affinity purification assay in a *Xenopus* kidney cell line ^17^. Here, in kidney-induced animal explants, we characterized a candidate from this approach, <u>s</u>ingle-<u>s</u>tranded DNA <u>b</u>inding <u>p</u>rotein <u>2</u> (Ssbp2), that was found to interact with the Ldb1-Lhx1 complex in kidney cells of both origins prompting us to investigate its function in this tissue.

Ssbp genes are evolutionarily conserved from *Drosophila* to humans ^21, 22^. Structurally, all three Ssbp vertebrate’s proteins (Ssbp2-4) contain a proline-rich transactivation domain ^23^ and a LUFS domain required for nuclear localization, homotetramerization and interaction with Ldb proteins ^24, 25^. Ssbp proteins bind and stabilize Ldb proteins from proteasomal degradation, therefore promoting their transcriptional function in multiple developmental processes, including wing development in *Drosophila*, axis formation in *Xenopus* and head morphogenesis in mice and *Xenopus* ^22, 25–27^. In mammals, Ssbp2 acts as tumor suppressor protein in several cancers including leukemia ^27^, pancreatic cancer ^28^, prostate cancer ^29^, oligodendroglioma ^30^, and esophageal squamous cell carcinoma ^31^, highlighting roles in malignant transformation. Ssbp2 knockout mice are born at lower frequency than expected and die prematurely. In addition to formation of carcinomas and lymphomas in multiple organs they develop glomerular nephropathy ^27^. However, Ssbp2 roles in kidney development have not been explored.

Here, we demonstrate the importance of Ssbp2 in pronephric kidney development of *Xenopus laevis*. In this model system, *ssbp2* is expressed in the kidney anlage, glomus and pronephric tubule. *Ssbp2* knockdown experiments demonstrate defective tubule and glomus morphogenesis, revealing a previously undescribed function in kidney organogenesis. Our results contribute to show a fundamental function for the Ldb1-Lhx1-Ssbp transcriptional complexes in multiple tissues during development.

## Results

### Ssbp2 interacts with Ldb1-Lhx1 in kidney-induced animal cap explants

To identify candidate partners associated with the constitutive-active transcriptional complex Ldb1-Lhx1 (LLCA) involved in nephrogenesis, we previously applied a tandem-affinity purification (TAP) approach using *Xenopus* renal A6 epithelial kidney cells ^17^. Here we employed an *in vitro* system inducing pronephric tissues by treatment of *Xenopus* animal cap explants with retinoic acid (RA) and activin ^32–34^ and applied the same purification protocol ^17^. Following injection of 2-cell stage embryos with *TAP-LLCA* mRNA, animal caps were dissected, treated with activin and RA, and protein complexes isolated through TAP (Supp. Fig. S1a). We subjected samples to nano-liquid chromatography coupled with tandem mass spectrometry (nanoLC-MS/MS) and identified 42 proteins exclusively associated to TAP-LLCA (Supp. Fig. S1b). Among those proteins, seven are expressed during *Xenopus* pronephros development ^35^ (Supp. Fig. S1b). Interestingly, <u>s</u>ingle-<u>s</u>tranded DNA <u>b</u>inding <u>p</u>rotein 2 (Ssbp2) was also identified in our previously published experimental approach ^17^ (Supp. Fig. S1b). While association of Ssbp proteins and Ldb1-Lhx1 regulates axis and head development ^21, 36–39^, Ssbp2 function in this ternary core complex has not been reported in kidney organogenesis.

### Ssbp2 is expressed during development of the *Xenopus* pronephros

To investigate a role for Ssbp2 in *Xenopus* pronephros formation, we initially assessed its expression pattern by whole mount *in situ* hybridization (WISH). *ssbp2* transcripts were detected in the circumblastoporal region of the gastrula (Fig. 1a), in the eye field, neural folds, and prospective olfactory placode in early neurula (Fig. 1b). In late neurula, *ssbp2* is expressed in the neural tube, somites and in the intermediate and lateral plate mesoderm (Fig. 1c-e’). Expression in early tailbuds becomes restricted to the head, notochord, neural tube, somites and pronephric anlage (Fig. 1f-h’). At later stages, *ssbp2* is also present in the branchial arches and the ventral blood islands (Fig. 1i,l). In the pronephros, *ssbp2* mRNAs are detected in the glomus, and in the proximal and distal tubules (Fig. 1i-n’). These results indicate that *ssbp2* transcripts are present throughout *Xenopus* pronephric kidney development.

**Figure 1.**
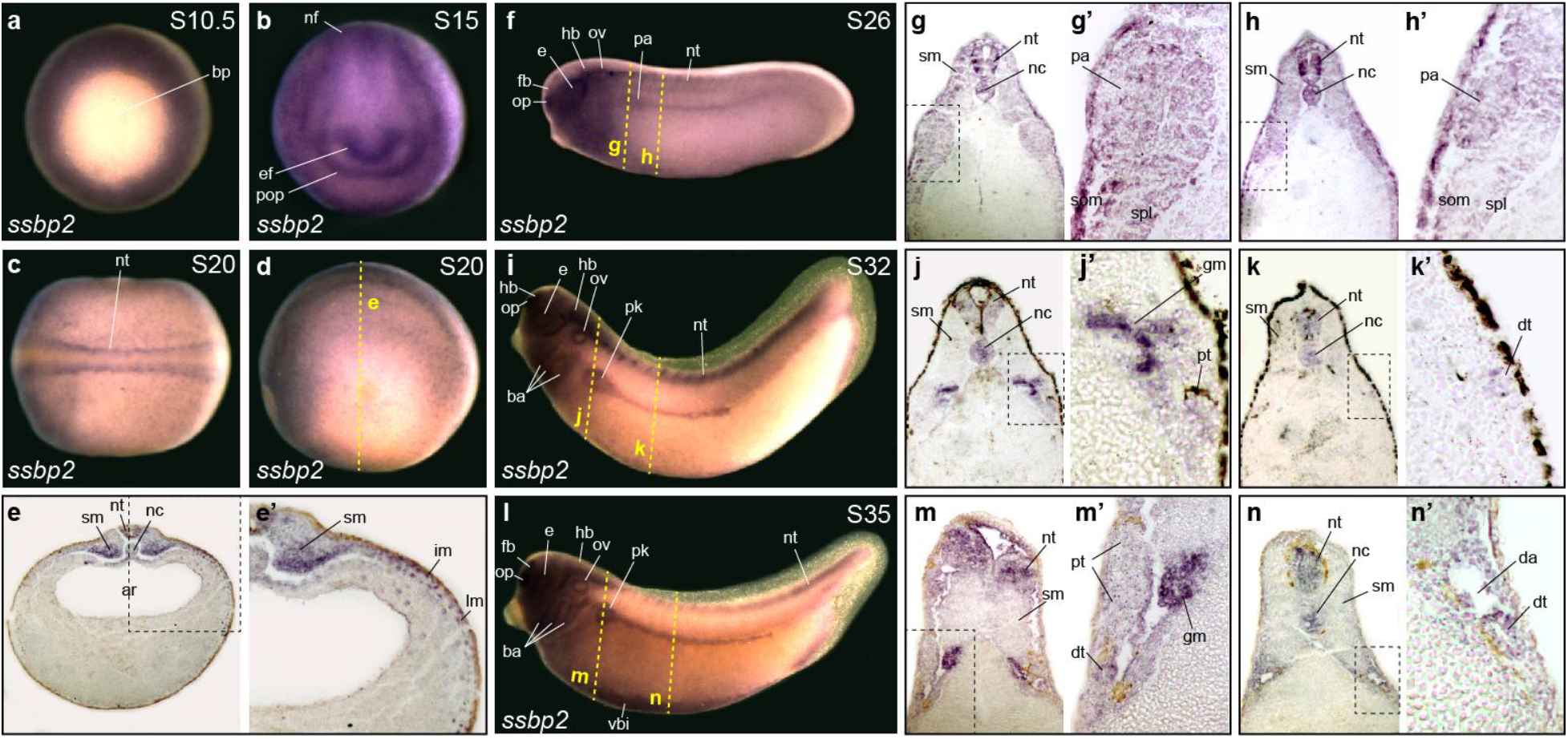
*ssbp2* is expressed in the pronephric anlage and the pronephric kidney during *Xenopus* development. *ssbp2* expression in *Xenopus* embryos by whole-mount *in situ* hybridization (WISH). (**a**) Early gastrula stage embryo (S10.5). Vegetal view. bp: blastopore. (**b**) Early neurula stage embryo (S15). Anterior view, dorsal up. ef: eye field; pop: presumptive olfactory placode; nf: neural folds. (**c,d**) Late neurula stage embryo (S20). (**c**) Dorsal view, anterior to the left. nt: neural tube. (**d**) Lateral view, anterior to the left, dorsal up. (**e**) Histological preparation (transverse section) of the embryo showed in *d* (dotted line). sm: somitic mesoderm; nc: notochord; ar: archenteron. (**e’**) Magnification of the dotted square shown in *e*. im: intermediate mesoderm; lm: lateral plate mesoderm. (**f**) Early tailbud stage embryo (S26). Lateral view. op: olfactory placode; fb: forebrain; e: eye; hb: hindbrain; ov: otic vesicle; pa: pronephric anlage. (**g,h**) Histological preparations (transverse sections) of the embryo shown in *f* (dotted lines). (**g’,h’**) Magnification of the dotted squares shown in *g* and *h,* respectively. som: somatic layer of the lateral mesoderm; spl: splanchnic layer of the lateral mesoderm. (**i**) Mid-tailbud stage embryo (S32). Lateral view. ba: brachial arches; pk: pronephric kidney. (**j,k**) Histological preparations (transverse sections) of the embryo shown in *i* (dotted lines). (**j’,k’**) Magnification of the dotted squares showed in *j* and *k,* respectively. gm: glomus; pt: proximal tubule; dt: distal tubule. (**l**) Late-tailbud stage embryo (S35). Lateral view. vbi: ventral blood islands. (**m,n**) Histological preparations (transverse sections) of the embryo shown in *l* (dotted lines). (**m’,n’**) Magnification of the dotted squares shown in *m* and *n,* respectively. da: dorsal aorta. Representative embryos are shown.

### Knockdown of Ssbp2 leads to abnormal pronephros

To test Ssbp2 function in kidney development, we performed loss-of-function experiments targeting both pseudo-alleles of *ssbp2* mRNA with a translation blocking morpholino oligonucleotide (*ssbp2*-MO) (Fig. 2a). In the absence of an antibody recognizing *Xenopus* Ssbp2 protein, we tested the efficacy of *ssbp2*-MO injecting 2-cell stage embryos with a fusion mRNA containing the full-length *ssbp2* cDNA in frame with *eGFP* (*ssbp2-eGFP*) (Supp. Fig. S2a). *ssbp2-eGFP* mRNA was injected alone or co-injected with a standard morpholino control (St-MO) or *ssbp2*-MO. While fluorescence was detected in *ssbp2-eGFP* and *ssbp2-eGFP* + St-MO injected embryos, it was not detected upon coinjection of *ssbp2-eGFP* + *ssbp2*-MO (Supp. Fig. S2b-e).

**Figure 2.**
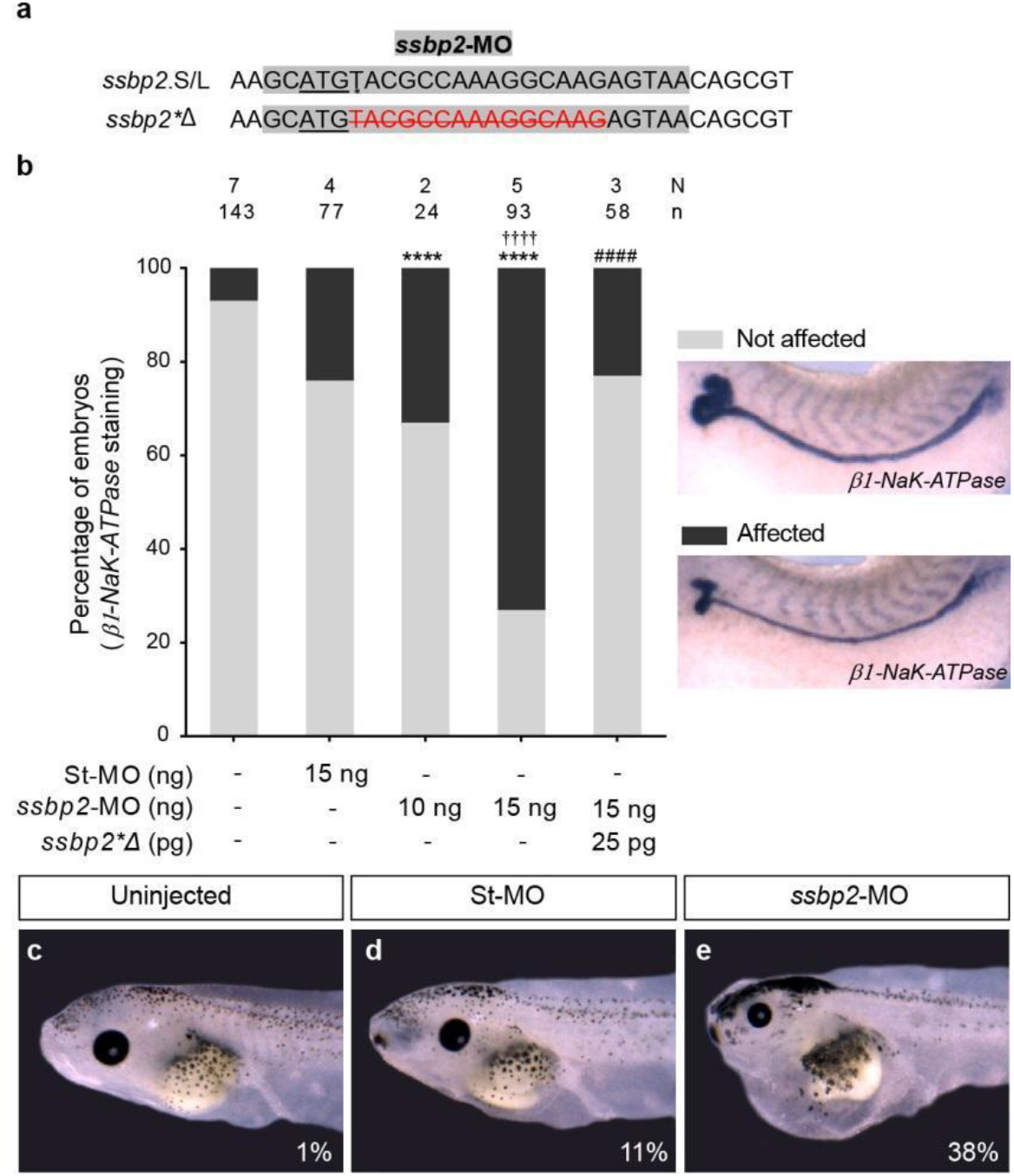
Ssbp2 morpholino knockdown affects pronephric kidney development and function. (**a**) *ssbp2*.S/L pseudo-alleles and *ssbp2*Δ* partial sequences. The initiation codon ATG is underlined. *ssbp2*-MO target site is highlighted in grey. Note that *ssbp2*Δ* possesses a 15 nucleotides deletion (crossed out red characters) to prevent *ssbp2*-MO binding. (**b**) 8-cell stage *Xenopus* embryos were injected into a single V2 blastomere as indicated, fixed at stage 39 and subjected to WISH for the *β1-NaK-ATPase* to assess formation of the pronephric tubule. The uninjected contralateral side was used as an internal control. The percentage of embryos showing affected pronephros was quantified. Data in the graph is presented as mean. Statistical significance was evaluated using *Chi-square* test (****p < 0.0001). * represent the comparison to the uninjected group, † represent the comparison to the St-MO injected group and # represent the comparison to the *ssbp2*-MO 15 ng injected group. (**c-e**) 4-cell stage *Xenopus* embryos were injected into both ventral blastomeres as indicated and edemas formation was analyzed at tadpole stage 45. (**c**) Uninjected embryo (1% with edema; n = 88; N = 3). (**d**) St-MO 30 ng injected embryo (11% with edema; n = 81; N = 3). (**e**) *ssbp2*-MO 30 ng injected embryos (38% with edema; n = 68; N = 3). N: number of independent experiments, n: number of embryos. Representative embryos are shown.

We evaluated the effect of *ssbp2* knockdown in pronephros formation injecting *ssbp2*-MO in one V2 blastomere of 8-cell stage embryos (1 x V2) ^40^ targeting one nephron and allowing us to compare injected and uninjected sides of the same embryo as the scoring system. We assessed *β1-NaK-ATPase* expression (marker of differentiated pronephric tubule) by WISH and observed a dose-dependent increase in the number of affected embryos (Fig. 2b). Injecting 15 ng of *ssbp2*-MO resulted in 74% of the embryos with affected pronephros without evidence of toxicity and so selected as experimental dose. By contrast, injecting 15 ng of St-MO resulted in significantly fewer affected embryos (Fig. 2b). The phenotype specificity was verified by coinjecting *ssbp2*-MO and *ssbp2*\*Δ mRNA. The *ssbp2*\*Δ mRNA has a deletion of 15 nucleotides following *ssbp2* first ATG, preventing binding of the morpholino (Fig. 2a). Importantly, *ssbp2*\*Δ mRNA coinjection significantly rescued *β1-NaK-ATPase* expression pattern (Fig. 2b).

To assess kidney function, embryos were injected with *ssbp2*-MO or St-MO in both ventral blastomeres of 4-cell stage embryos ^41^, allowed to develop until stage 45 and analyzed for edema formation. Uninjected and St-MO injected tadpoles presented low incidences of edema formation (1% and 11%, respectively), whereas 38% of *ssbp2*-MO injected tadpoles developed edema (Fig. 2c-e). Together, these experiments suggest that Ssbp2 loss-of-function results in defective pronephros formation and altered kidney function.

### Ssbp2 is not essential for establishment of the pronephric field

In *Xenopus*, *lhx1* knockdown impairs specification of the entire kidney field affecting expression of genes from glomus, proximal and distal kidney segments ^16^. To understand Ssbp2 role in pronephric development, we analyzed by WISH *pax8* expression in the early pronephric field. *Pax8* is a transcription factor essential for the earliest steps of vertebrate pronephric development and together with *lhx1* are among the earliest genes expressed in the kidney primordium ^42, 43^. Injection of *ssbp2*-MO did not alter *pax8* expression domain relative to the control uninjected side (Supp. Fig. S3a-c). We also evaluated *osr2* (odd-skipped related transcription factor 2) expression in the pronephric field as its expression precedes activation of the early pronephric markers *pax8* and *lhx1* ^42, 44^. The expression domain *of ors2* was not affected in most *ssbp2*-MO injected embryos (Supp. Fig. S3d-e), indicating that Ssbp2 function is not essential for renal progenitor cell field specification.

### Ssbp2 is required for glomus development

Expression of *ssbp2* was detected in the glomus at tailbud stages (Fig. 1). As readout of glomus differentiation, we analyzed *ssbp2* depleted embryos for the podocyte differentiation marker *nphs1* (nephrin) at stage 35, before podocytes function begins ^45^ (Fig. 3a-h). The *nphs1* expression domain showed a slight reduction in the *ssbp2*-MO injected side (Fig. 3f,g,h), whereas no differences were observed between injected and uninjected sides of St-MO treated embryos (Fig. 3c,d,e). As the glomus is the deepest structure of the pronephros, histological transverse sections were prepared to assess in more detail its morphology (Fig 3b,e,h). While the structure of the glomus was similar in kidneys of control embryos (Fig. 3b,e), the surface area occupied by podocytes was reduced in embryos unilaterally injected with *ssbp2*-MO (Fig. 3h). This result indicates that in the pronephros, Ssbp2 is necessary for normal glomus development.

**Figure 3.**
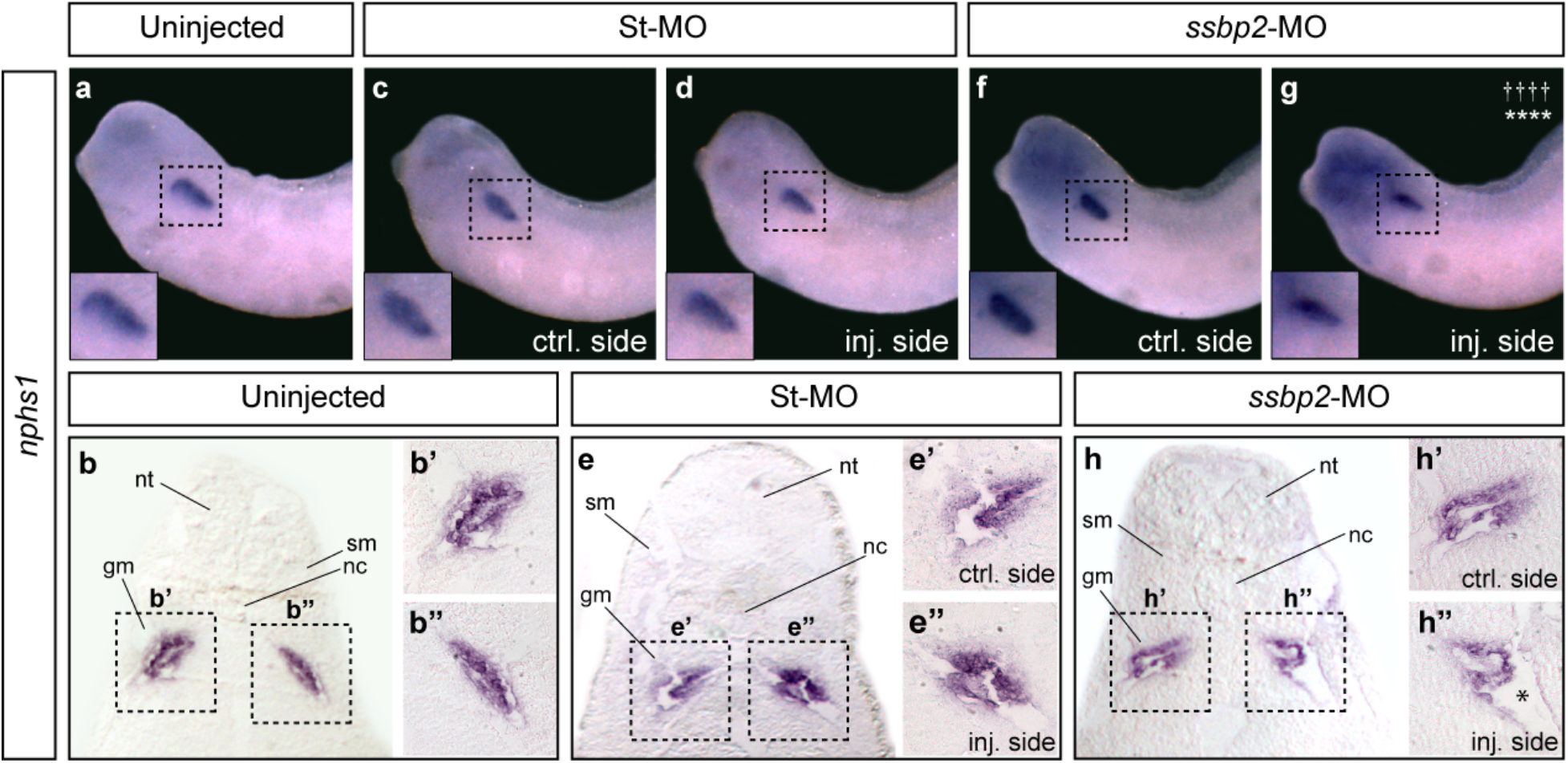
Ssbp2 is required for normal glomus development. 8-cell stage *Xenopus* embryos were injected into a single V2 blastomere as indicated and the expression of *nphs1* in the glomus was analyzed by WISH at stage 35. The uninjected contralateral side was used as an internal control. (**a,b**) Uninjected embryo (5% affected, n = 37, N = 2). (**c-e**) St-MO 15 ng injected embryo (14% affected, n = 37, N = 2). (**f-h**) *ssbp2*-MO 15 ng injected embryo (46% affected, n = 40, N = 2). Statistical significance was evaluated using *Chi-square* test (****p < 0.0001). * represent the comparison to the uninjected group and † represent the comparison to the St-MO injected. N: number of independent experiments, n: number of embryos. (**b,e,h**) Histological preparations (transverse sections). nt: neural tube; sm: somites; nc: notochord; gm: glomus. (**b’,b’’,e’,e’’,h’,h’’**) Magnification of the glomus enclosed by the black squares in *b, e* and *h*. * indicates a reduction in the surface of the developing glomerular filtration barrier. Representative embryos are shown.

### Ssbp2 depletion impairs tubule morphogenesis without affecting somites development

*Ssbp2* is expressed in the paraxial mesoderm (Fig. 1) and interactions between developing somites and pronephros influence patterning of both tissues ^16, 46–50^. Therefore, we asked whether the pronephric phenotype observed in *ssbp2*-depleted embryos is secondary to somites developmental defects. Immunostaining with the muscle-specific antibody 12/101 ^51^ showed no differences in somite morphology between controls (uninjected and St-MO injected) and *ssbp2*-MO injected embryos (Fig. 4a-f), suggesting that defects associated with Ssbp2 loss-of-function in the pronephros are not the result of alterations in paraxial mesoderm development.

**Figure 4.**
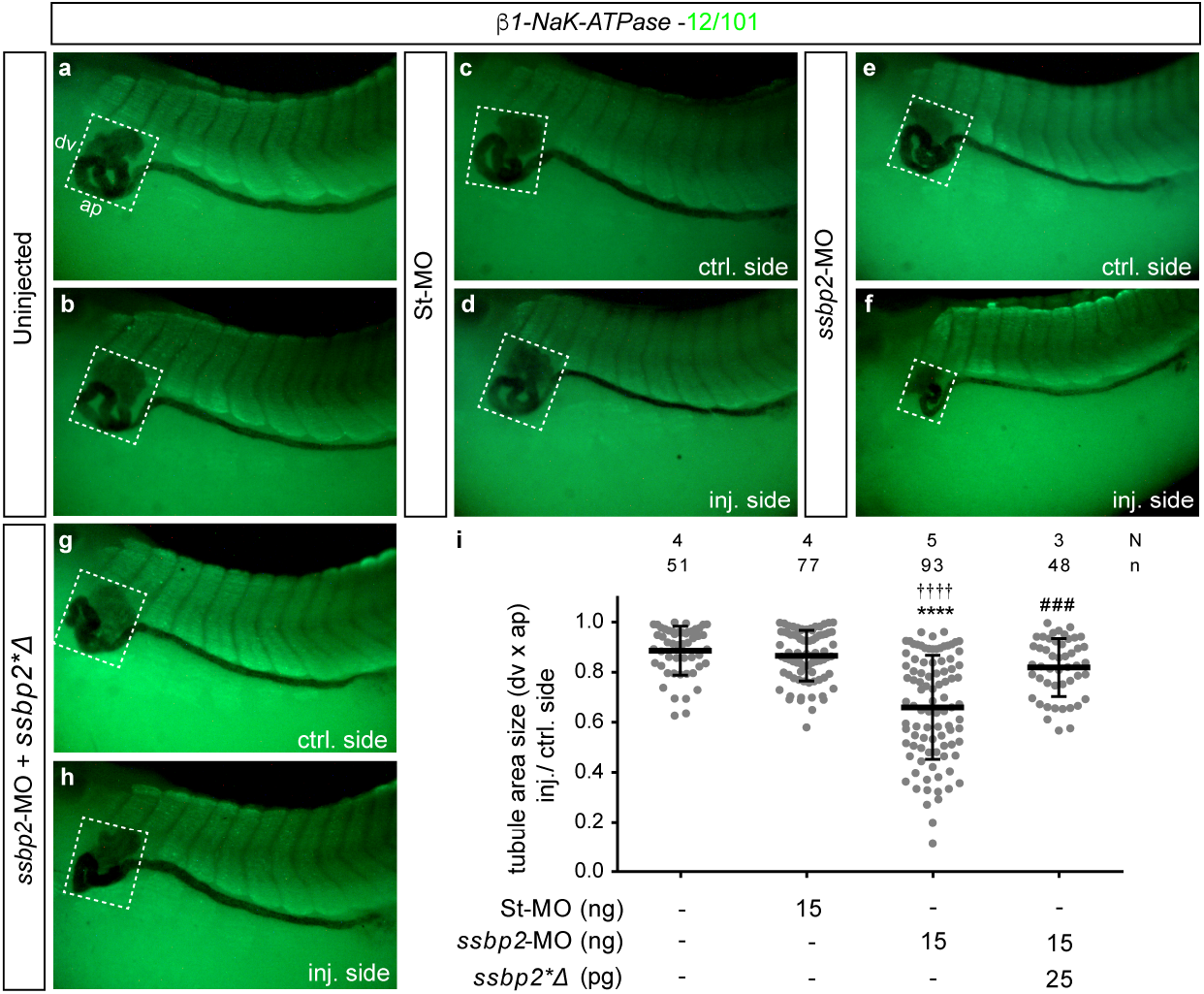
Ssbp2 loss-of-function impairs tubule morphogenesis. 8-cell stage *Xenopus* embryos were injected into a single V2 blastomere as indicated, fixed at stage 39 and subjected to WISH for *β1-NaK-ATPase* followed by immunostaining with 12/101 antibody to assess pronephros and somites development, respectively. The uninjected contralateral side was used as the internal control. (**a,b**) Uninjected embryo. (**c,d**) St-MO 15 ng injected embryo. (**e,f**) *ssbp2*-MO 15 ng injected embryo. (**g,h**) Rescue experiment. *ssbp2*-MO 15 ng + *ssbp2*Δ* mRNA 25 pg coinjected embryo. The white square encloses the tubule convoluted area. dv: dorsal-ventral; ap: anterior-posterior. Lateral views, anterior to the left. Representative embryos are shown. (**k**) Quantification of the tubule convoluted area (dv x ap). The ratio between the injected and control side is shown. Data in graph is presented as mean and standard deviation. Each point represents a single embryo. Statistical significance was evaluated using *Kruskal–Wallis* test and *Dunn’s* multiple comparisons test (****p < 0.0001; ***p < 0.001). * represent the comparison to the uninjected group, † represent the comparison to the St-MO injected group and # represent the comparison to the *ssbp2*-MO injected group. N: number of independent experiments, n: number of embryos.

To characterize the pronephric tubule defects associated with Ssbp2 loss-of-function, we performed a quantification of the total tubular convoluted surface in embryos subjected to *β1-NaK-ATPase* WISH (Fig. 4). We established that *ssbp2* knockdown led to less convoluted proximal and distal tubule resulting in a significant reduction of the tubular surface (Fig. 4e,f,i). Importantly, this phenotype was rescued by *ssbp2*\*Δ mRNA coinjection (Fig. 4g-i) demonstrating *ssbp2*-MO specificity.

Next, we investigated the tubulogenesis defect by analyzing expression of specific tubular segment markers. In *Xenopus*, the pronephric tubule branches at its most proximal end to generate three ciliated structures called nephrostomes that receive fluid derived from the coelom ^50^. As shown with *pax2* expression, nephrostomes do not properly form and remain jointly in *ssbp2* knockdown embryos (Fig. 5a-e). Similarly, proximal tubules assessed by *slc5a1* (sodium/glucose cotransporter) expression appeared smaller in size relative to the uninjected side and to St-MO injected embryos (Fig. 5f-j). Additionally, we looked in tailbud stages at *pax8* as expression of this gene becomes restricted to the pronephric tubule during kidney morphogenesis. We observed a reduced expression domain of *pax8* in the proximal tubule upon *ssbp2* depletion (Supp. Fig. S4d,e), while significantly fewer embryos were affected in uninjected and St-MO injected groups (Supp.Fig. S4a-c). Analysis of distal tubule development by WISH against *clcnkb* (chloride channel) established that *ssbp2* knockdown led to reduced tubule size (Fig 5k-o). To confirm this observation, we measured the length of *hoxb7* (homeobox B7) expression domain in the distal tubule of stage 32 embryos. The evaluation revealed that this tubule segment was significantly shorter on the *ssbp2*-depleted side relative to the control side of the same embryo (Supp. Fig. S4f-k), while the connecting tubule appeared unaffected (Supp. Fig. S4a-j). Taken together, these results show that while all components of the tubule are present in depleted embryos, Ssbp2 function is required for normal morphogenesis of the proximal and distal tubule segments.

**Figure 5.**
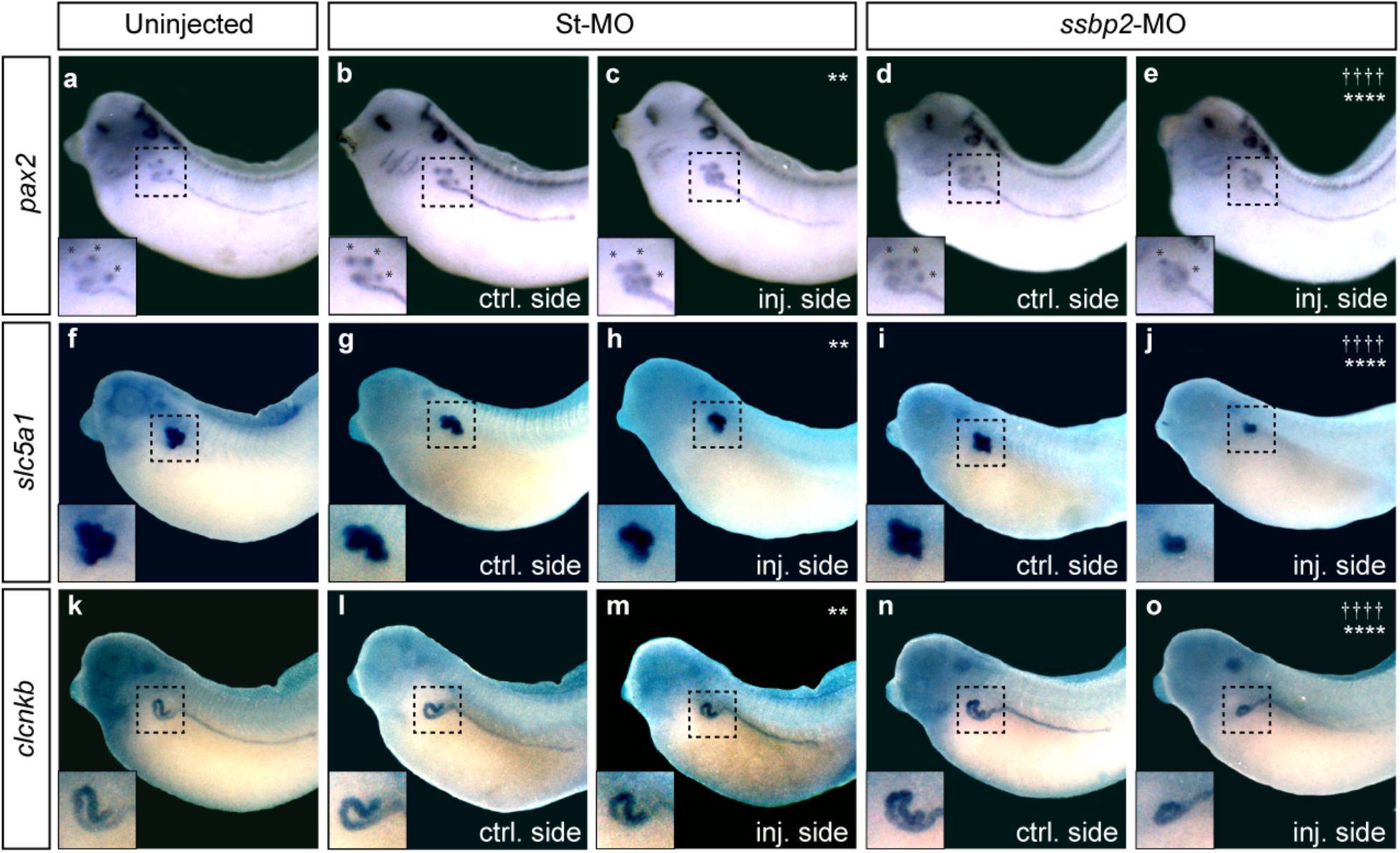
Ssbp2 depletion reduces the expression domain of nephrostomes, proximal and distal tubule markers. 8-cell stage *Xenopus* embryos were injected into a single V2 blastomere as indicated. The uninjected contralateral side was used as an internal control. (**a-e**) WISH for *pax2* in stage 32 embryos. * indicate the position of the nephrostomes. (**a**) Uninjected embryo (8% affected, n = 37, N = 3). (**b,c**) St-MO 15 ng injected embryo (23% affected, n = 40, N = 2). (**d,e**) *ssbp2*-MO 15 ng injected embryo (66% affected, n = 51, N = 3). (**f-j**) WISH for *slc5a1* in stage 39 embryos. (**f**) Uninjected embryo (4% affected, n = 47, N = 2). (**g,h**) St-MO 15 ng injected embryo (19% affected, n = 39, N = 2). (**i,j**) *ssbp2*-MO 15 ng injected embryo (50% affected, n = 34, N = 2). (**k-o**) WISH for c*lcnkb* in stage 39 embryos. (**k**) Uninjected embryo (3% affected, n = 35, N = 2). (**l,m**) St-MO 15 ng injected embryo (23% affected, n = 41, N = 2). (**n,o**) *ssbp2*-MO 15 ng injected embryo (49% affected, n = 33, N = 2). Magnifications of the pronephric tubules enclosed by the black squares are shown in the left-bottom corner. Statistical significance was evaluated using *Chi-square* test (****p < 0.0001; **p < 0.01). *, † represent the comparison to the uninjected and the St-MO injected groups, respectively. Representative embryos are shown.

### Depletion of Ssbp2 impairs pronephros terminal differentiation

Immunostaining with 3G8 and 4A6 antibodies were conducted to assess differentiated proximal tubules (stage 37/38 embryos) and distal and connecting tubules (stage 42 embryos), respectively ^50^ (Fig. 6). 3G8 staining revealed a reduced proximal convoluted tubule in the *ssbp2-*MO injected side relative to the uninjected side of the embryo (Fig. 6e-f) and relative to control treatments (Fig. 6a-d). To quantify the extent of pronephric proximal tubule development we calculated the pronephric index (PNI) ^52^. The PNI value is expressed as the number of tubule components on the uninjected side (left, Fig. 6a’,c’,e’) minus the number on the injected side (right, Fig. 6b’,d’,f’) of the embryo. Thus, significant reduction of the tubule components by experimental treatments results in high PNI values whereas treatments causing little, or no effect result in low numbers (Fig. 6g). The PNI scoring was significantly greater in *ssbp2*-depleted embryos than in St-MO and uninjected embryos (Fig. 6g). Further, 3G8 staining revealed that the proximal tubules of depleted embryos had shorter branches than St-MO and uninjected animals (Fig. 6a-f). Staining of distal and connecting tubules with 4A6 antibody also showed a reduction in distal tubule size in the *ssbp2*-MO injected side relative to the uninjected side (Fig. 6l,m) and control treatments (Fig. 6h-k). Histological sections confirmed reduced staining and convolution of the distal tubule because of *ssbp2* knockdown (Fig. 6n), while the connecting tubules appear unaffected (Fig. 6o). Taken together and in agreement with our previous evidence (Figs. 4, 5 and Supp. Fig. S4), Ssbp2 loss-of-function impairs proximal and distal tubule terminal differentiation.

**Figure 6.**
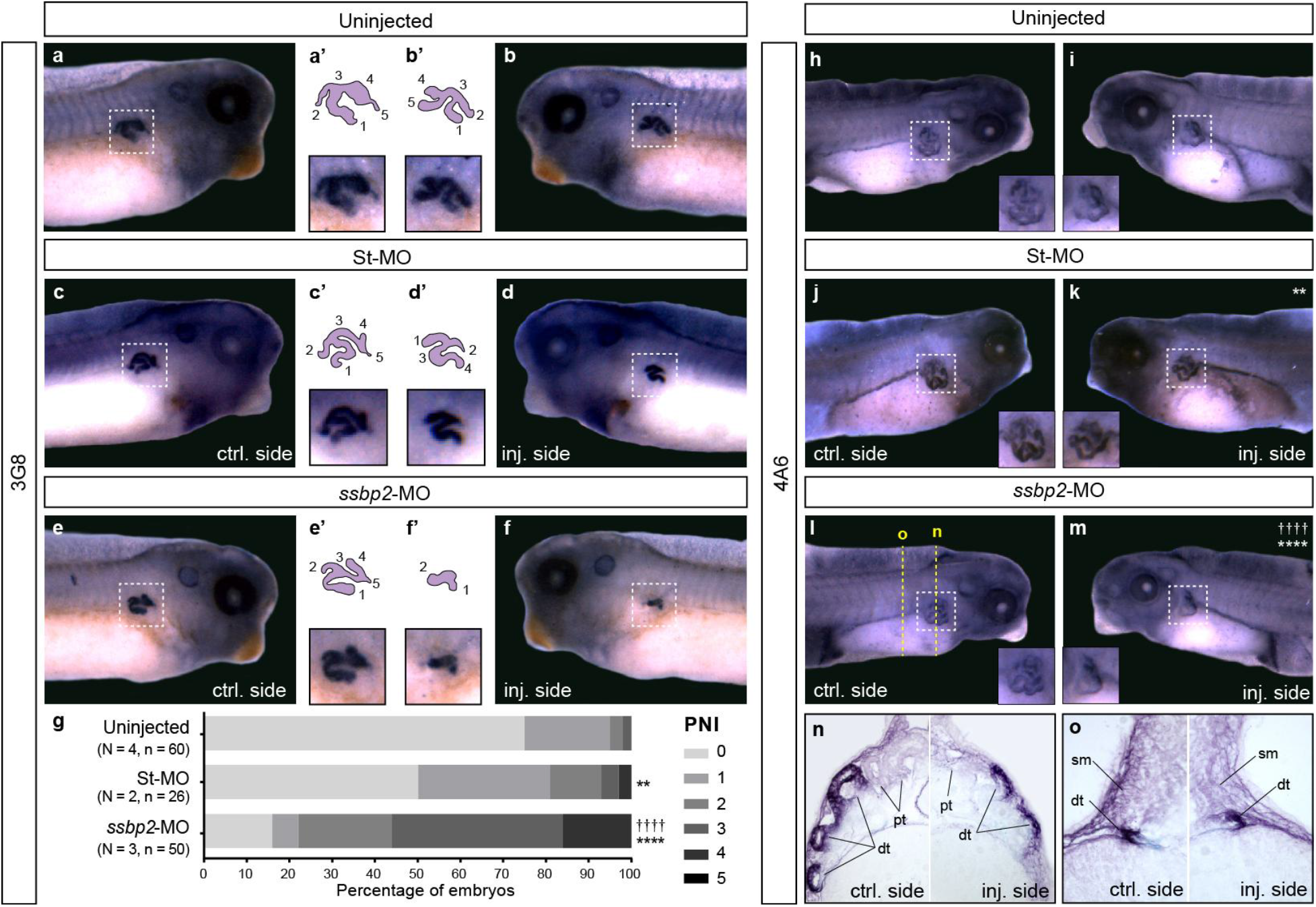
Ssbp2 depletion affects pronephros terminal differentiation. 8-cell stage *Xenopus* embryos were injected into a single V2 blastomere as indicated. The uninjected contralateral side was used as an internal control. (**a-f**) 3G8 whole-mount immunostaining was carried out at stage 37/38. (**a,b**) Uninjected embryo. (**c,d**) St-MO 15 ng injected embryo. (**e,f**) *ssbp2*-MO 15 ng injected embryo. (**a’-f’**) Magnifications (bottom) and schemes (top) of the proximal tubules enclosed by the white squares in *a-f.* Numbers indicate the tubule components. (**g**) The pronephric index (PNI) was scored as the difference between the number of proximal tubules components on both sides of the same embryo (PNI = 0 indicates two identical proximal tubules). The percentage of embryos exhibiting different PNI values is shown. Data in the graph is presented as mean. (**h-o**) 4A6 whole-mount immunostaining were carried out at stage 42. Magnifications of the distal tubules enclosed by the white square are shown for each side of the embryo. (**h,i**) Uninjected embryo (8% affected; n = 40; N = 2). (**j,k**) St-MO 15 ng injected embryo (28% affected; n = 34; N = 2). (**l,m**) *ssbp2*-MO-MO 15 ng injected embryo (59% affected; n = 61; N = 3). Statistical significance was evaluated using *Chi-square* test (****p < 0.0001; **p < 0.01). *, † represent the comparison to the uninjected and the St-MO injected groups, respectively. N: number of independent experiments, n: number of embryos. (**n, o**) Histological preparations (transverse sections) of the embryo shown in *l* and *m*. pt: proximal tubule; sm: somites; dt: distal tubule. Representative embryos are shown.

## Discussion

Understanding the molecular mechanisms responsible for kidney specification and morphogenesis is fundamental to comprehend the etiology of renal diseases. In this study our goal was to identify interacting partners of the Ldb1-Lhx1 transcriptional complex, with a conserved and essential role in nephric development. The tetrameric Ldb1-Lhx1 complex is required for cell fate acquisition, embryonic patterning, and organ development. Regulation of its transcriptional function and expression of target genes is thought to be modulated by accessory proteins ^53^. Here, we use a simple and efficient kidney induction assay ^32–34, 54, 55^ to identify binding partners of the Ldb1-Lhx1 complex employing our previously published TAP assay followed by LC-MS/MS ^17^. Among the identified proteins, several are expressed in the *Xenopus* pronephric kidney (Tubal3.1, Tubb4a, Hspd1, Hspa1b, Slc25a11, Etfa, Prdx1) ^35^. Validation and functional analysis of these candidate interactors might uncover new proteins with roles in kidney organogenesis. We focused this study on Ssbp2 since we identified it as a binding partner in *Xenopus* renal epithelial cells ^17^ and here in the *ex vivo* kidney-induction assay. Additionally, Ssbp proteins have been found to physically interact with Ldb1 ^21^. This interaction involves the LUFS domain of Ssbp protein and the LCCD domain (Ldb1-Chip conserved domain) of Ldb1 ^22^. Since the TAP-LLCA construct contains the LCCD domain in Ldb1 fragment, we speculate Ssbp2 interacts with constitutive-active Ldb1-Lhx1 through this region. Previously, Ssbp2 was found to bind Ldb1 in a yeast two-hybrid screening and with complexes containing Lhx1 in a mouse carcinoma cell line ^22, 36^. Our TAP results provide further evidence of this physical interaction.

The ability of Ssbp proteins to stabilized Ldb1 and enhance the transactivating function of the Ldb1-Lhx1 complex has been well documented ^21, 29, 37 19, 25, 33, 54^ and genetic interactions between Ldb, (LIM binding domain), LIM and Ssbp proteins have been previously shown in *Drosophila* ^21, 22^, *Xenopus* ^21^ and mice ^37, 38^. Critical to this complex formation is the maintenance of protein stoichiometry. In fact, excess of free Ldb1 (not associated with Lhx1) is targeted for proteasome degradation ^56^. Ssbp functions suppressing this degradation, maintaining the stoichiometry between Ldb1, Lhx1 and Ssbp ^26^. Based on our loss-of-function experiments, we can speculate that Ssbp2 knockdown is directly altering the complex stoichiometry and indirectly reducing Ldb1 protein abundance, perturbing the complex assembly. Future efforts will focus on uncovering Ssbp2 molecular mechanism and how it is related to the Ldb1-Lhx1 complex regulation.

We describe for the first time the ssbp2 expression pattern in developing *Xenopus* embryos. In agreement with Ssbp2 knockout mice occasionally displaying smaller heads ^57^, in *Xenopus* embryos we detected *ssbp2* transcripts in dorsal midline tissues and head structures. The evidence suggests a role of Ssbp2 in development of anterior-head structures and a possible functional redundancy with Ssbp3 in these tissues ^57^. Importantly, we detected by WISH the presence of *ssbp2* mRNA transcripts in the intermediate mesoderm and later in the glomus and tubule of the developing pronephric kidney. Our *ssbp2* expression results agree with a recent single-cell sequencing analysis of *Xenopus* tadpole pronephros, where this mRNA was found to be enriched in the glomus ^2^. Moreover, our results and the identification of SSBP2 as a human glomerular-enriched gene ^58^ argue in favor of this protein having an evolutionarily conserved role in glomerulus/glomus development and function.

We demonstrate that *ssbp2* targeted knockdown in blastomeres contributing to pronephric tissues leads to defects in kidney development. Importantly, rescue experiments coinjecting *ssbp2* mRNA, prove that the observed phenotype is specific to Ssbp2 loss-of-function. While our evidence points to the fact that this protein is not essential for pronephric field specification, abnormal glomus formation in depleted embryos is consistent with the finding that Ssbp2 null mice exhibit chronic glomerular nephropathy ^27^. Nevertheless, dissecting Ssbp2 function in the glomus formation will require identification of its role in the podocytes transcriptional network.

Interestingly, proximal tubules branching was impaired upon *ssbp2* depletion as well as normal elongation of the distal tubule. Staining of differentiated proximal (3G8) and distal (4A6) regions of the tubule revealed reduced morphogenesis and differentiation of these segments. While *lhx1* knockdown impairs specification of the entire kidney field dramatically reducing expression of nephrogenic factors, *ssbp2* depletion has only mild effects on early pronephric field marker genes. Given that unlike *lhx1* ^15, 16^, *ssbp2* mRNAs were not detected in the pronephric anlage at the time renal progenitors are specified; this result is not unexpected. Differences between *lhx1* ^16^ and *ssbp2* knockdown embryos might be due to the requirement of different binding partners by Ldb1-Lhx1 throughout kidney development. In this regard, we have described a functional interaction between LLCA and Fry protein, regulating miRNAs levels and driving kidney field specification. Evidence from other Ssbp-Ldb1-Lhx1 complexes in *Xenopus* ^59^ and mice ^60^ support the idea that this core complex function is driven by different binding partners likely temporally and spatially restricted, recruited for control of specific gene networks and developmental programs. However, another possibility is that other Ssbp family proteins exert a compensatory role reducing the effect of Ssbp2 depletion, this hypothesis will require further investigation.

Molecular interactions between Ssbp and Ldb proteins are evolutionarily ancient ^22, 61^ and supply a fundamental function in transcription regulation, constituting a complex widely used among eukaryotes with roles in cell differentiation, cell identity, and tissue-specific gene regulation. Here we show that Ssbp2, a member of the Ssbp family, physically interacts with constitutive-active Ldb1-Lhx1 in *Xenopus* kidney-induced cells. With functional studies we demonstrate its functional requirement for normal pronephric kidney morphogenesis and differentiation.

## Materials and methods

### Ethics statement

This study was carried out in strict accordance with the recommendations in the Guide for the Care and Use of Laboratory Animals of the NIH and the ARRIVE guidelines (https://arriveguidelines.org/). The animal care protocol was approved by the Comisión Institucional para el Cuidado y Uso de Animales de Laboratorio (CICUAL) of the School of Applied and Natural Sciences, University of Buenos Aires, Argentina (Protocol #64).

### *Xenopus* embryos preparation

*Xenopus laevis* embryos were obtained by natural mating. Adult frog’s reproductive behavior was induced by injection of human chorionic gonadotropin hormone. Eggs were collected, de-jellied in 3% cysteine (pH 8.0), maintained in 0.1X Marc’s Modified Ringer’s (MMR) solution and staged according to Nieuwkoop and Faber ^62^. The embryos were placed in 3% ficoll prepared in 1X MMR for microinjection.

### Animal caps and Tandem Affinity Purification (TAP)

2-cell stage embryos were injected into the animal pole of both blastomeres with 1 ng of *TAP.LLCA* mRNA and allowed to develop until stage 8. To induce explanted animal caps into pronephric tissue, dissected animal caps from injected and uninjected sibling embryos were cultured in 1X Steinberg’s solution for 6 hours ^53^ in the presence of 10 ng/ml activin (Sigma, A4941) and 1X 10^-4^ M retinoic acid (RA) (Sigma, R2625), then washed in 1X Steinberg’s solution, and snap-freeze for later use. Approximately 300 animal caps from 7 independent experiments were collected and processed for TAP following manufacturer’s instructions (InterPlay Mammalian TAP System, Agilent Technologies). Samples subjected to TAP were sent for identification of proteins by nanoLC/MS/MS to MS Bioworks, LLC (Ann Arbor, MI) and processed as previously described ^17^. The results were analyzed by the Scaffold software.

### Plasmid constructs for mRNA synthesis

pCS2+.TAP.LLCA construct has been previously described ^17^. pCS2+.ssbp2.eGFP was made by PCR amplification of full-length ssbp2 cDNA from pCMV.Sport6.ssbp2 (Dharmacon) plasmid and cloned in frame 5’ of the eGFP into pCS2+.eGFP digested with *Nco*I and *BamH*I. For rescue experiments we made the pCS2+.ssbp2*Δ construct that lacks 15 nucleotides following the ATG. This construct was made by PCR amplification from pCMV.Sport6.ssbp2 and cloned into pCS2+ as a *BamH*I/*EcoR*I fragment. Constructs were verified by sequencing.

Primers for cloning ssbp2*Δ into pCS2+

ssbp2*Δ: 5’-AAGGATCCATGAGTAACAGCGTACC-3’

ssbp2.R: 5’-AAAGAATTCTCACACGCTCATTGTCATGCTAGG-3’

Primers for cloning ssbp2 cDNA into pCS2+.eGFP

ssbp2GFP.F: 5’-AAAGGATCCAGCATGTACGCCAAAGGCAAG-3’

ssbp2GFP.R: 5’-AAACCATGGCCACGCTCATTGTCATGCTAGG-3’

### mRNA and morpholinos microinjections

Capped mRNA for *ssbp2*-GFP and *ssbp2*Δ* were transcribed *in vitro* using the AmpliCap SP6 High Yield Message Marker Kit (Cellscript) following linearization with *Hind*III and *Not*I, respectively. For Ssbp2-depletion studies, an Ssbp2 morpholino oligonucleotide (*ssbp2*-MO) (Gene Tools, LLC; 5’-TTACTCTTGCCTTTGGCGTACATGC-3’) that prevents the translation of both *ssbp2* pseudo alleles (*ssbp2*.S/L) mRNAs was used. A Morpholino Standard Control oligo (St-MO) was used as a negative control (Gene Tools, LLC; 5’-CCTCTTACCTCAGTTACAATTTATA-3’). To confirm the specificity of *ssbp2*-MO, both blastomeres of 2-cell stage embryos were injected into the animal pole with 1 ng *ssbp2-eGFP* mRNA in the presence or absence of St-MO or *ssbp2*-MO. Embryos were screened for green fluorescence at stage 10.5. To study pronephros development, 8-cell stage embryos were injected into one of the V2 blastomeres (1x V2) ^40^. To evaluate edema formation 4-cell stage embryos were injected into both ventral blastomeres (2x V). Embryos were injected with 10-30 ng of *ssbp2*-MO, 15-30 ng of St-MO and 25 pg of *ssbp2*Δ* mRNA per embryo.

### In situ hybridization and immunostaining

Whole-mount *in situ* hybridization was carried out as previously described ^63^. *Pax8* (gift from Tom Carroll), *β1-NaK-ATPase*, *nphs1* (gift from Oliver Wessely), *slc5a1* and *clcnkb* (gift from Rachel Miller) were linearized as previously described ^16, 64, 65^. *Ssbp2, hoxb7* and *pax2* (Dharmacon) were linearized with *Sal*I. *Osr2* (gift from José Luis Gómez-Skarmeta) was linearized with *EcoR*I. All linearized constructs were transcribed with T7 for antisense probe synthesis. For whole-mount immunostaining with 12/101 (DSHB Cat# 12/101, RRID:AB_531892), 3G8 (EXRC Cat# 3G8.2C11, RRID:AB_10013600) and 4A6 (EXRC Cat# 4A6.2C10) we followed the protocol previously described ^16^. Goat anti-mouse Alexa Fluor 488 (Invitrogen, EXRC: AB_2534069) was used as secondary antibody for 12/101. For preparation of histological slides, we followed the protocol previously described ^66^. Xenbase (http://www.xenbase.org/, RRID:SCR_003280) was used as a source of information on gene expression, sequences, developmental stages, and anatomy.

### Image analysis

Images of fixed whole embryos were collected with a Leica DFC420 camera attached to a Leica L2 stereoscope. Histological slides were imaged using a digital camera (Infinity 1; Lumera Corporation) attached to a light-field microscope (CX31: Olympus). GFP fluorescence was visualized under a Discovery V8 (Zeiss) stereomicroscope and imaged with a MicroPublisher 3.3 RTV camera (Q Imaging). Pronephros morphometries were quantified from fixed embryos using ImageJ software (https://fiji.sc/). Pronephic index (PNI) ^52^ and the tubular area size ^67^ were calculated as previously described.

### Statistical analysis

Numbers of embryos (n) and independent experimental replicates (N) for animal studies are stated in graphs. Statistical analysis for Figures 2, 3, 5, S3 and S4 were performed using *Chi-square* tests (**p < 0.001, ***p < 0.001, ****p < 0.0001). Statistical analysis for Figures 4 and S4k were performed using *Kruskal–Wallis* tests and means between groups were compared using *Dunn’s multiple comparisons* tests (****p<0.0001, **p<0.01). For all statistical analyses we used Prism6, GraphPad Software, Inc.

## Acknowledgements

We would like to thank Oliver Wessely, Rachel Miller and José Luis Gómez-Skarmeta for kindly provide plasmid constructs; Eugenio Sewchuk for *Xenopus laevis* care; Lance Davidson for anti-mouse Alexa Fluor 488 antibody and RNA synthesis reagents. The laboratory of M. Cecilia Cirio received support for this work from the Agencia Nacional de Promoción Científica y Tecnológica of Argentina (PICT-2013 0381) and Consejo Nacional de Investigaciones Científicas y Técnicas (PIP-CONICET 11220150100577CO). The laboratory of Neil Hukriede received support for this work from the NIH 2R01 DK069403. ASC and IAL were supported by the CONICET Doctoral Fellowship Program.

## Author Contribution

A.S.C., M.G.C., I.A.L., D.H., N.A.H and M.C.C., conceived, designed, contributed, and analyzed data. A.S.C., M.G.C, I.A.L and M.C.C performed the *Xenopus* experiments. D.H. designed and performed the experiments for generating pCS2+.ssbp2.GFP and pCS2+.ssbp2*Δ constructs. The manuscript was written by M.C.C. and A.S.C with input from all authors.

## Data availability

All data generated or analyzed during this study are included in the published article.

## References

1. Zhou, X. & Vize, P. D. Proximo-distal specialization of epithelial transport processes within the Xenopus pronephric kidney tubules. Dev. Biol. 271, 322– 338 (2004).

2. Corkins, M. E. et al. A comparative study of cellular diversity between the Xenopus pronephric and mouse metanephric nephron. Kidney Int. 103, 77–86 (2023).

3. Torrey, T. W. Morphogenesis of the vertebrate kidney. Organogenesis 559–579 (1965).

4. Bouchard, M., Souabni, A., Mandler, M., Neubüser, A. & Busslinger, M. Nephric lineage specification by Pax2 and Pax8. Genes Dev. 16, 2958–2970 (2002).

5. Dressler, G. R. The Cellular Basis of Kidney Development. Annu. Rev. Cell Dev. Biol. (2006) doi:10.1146/annurev.cellbio.22.010305.104340.

6. Wingert, R. A. & Davidson, A. J. The zebrafish pronephros: a model to study nephron segmentation. Kidney Int. 73, 1120–1127 (2008).

7. Brändli, A. W. Towards a molecular anatomy of the Xenopus pronephric kidney. Int. J. Dev. Biol. 43, 381–395 (1999).

8. Hensey, C., Dolan, V. & Brady, H. R. The Xenopus pronephros as a model system for the study of kidney development and pathophysiology. Nephrol. Dial. Transplant. Off. Publ. Eur. Dial. Transpl. Assoc. - Eur. Ren. Assoc. 17 **Suppl 9**, 73–74 (2002).

9. Jones, E. A. Xenopus: a prince among models for pronephric kidney development. J. Am. Soc. Nephrol. 16, 313–321 (2005).

10. Blackburn, A. T. M. & Miller, R. K. Modeling congenital kidney diseases in Xenopus laevis. Dis. Model. Mech. 12, (2019).

11. Desgrange, A. & Cereghini, S. Nephron Patterning: Lessons from Xenopus, Zebrafish, and Mouse Studies. Cells 4, 483–499 (2015).

12. Lienkamp, S. S. Using Xenopus to study genetic kidney diseases. Semin. Cell Dev. Biol. 51, 117–124 (2016).

13. Barnes, J. D., Crosby, J. L., Jones, C. M., Wright, C. V & Hogan, B. L. Embryonic expression of Lim-1, the mouse homolog of Xenopus Xlim-1, suggests a role in lateral mesoderm differentiation and neurogenesis. Dev Biol 161, 168–178 (1994).

14. Chan, T. C., Takahashi, S. & Asashima, M. A role for Xlim-1 in pronephros development in Xenopus laevis. Dev. Biol. 228, 256–269 (2000).

15. Carroll, T. J. & Vize, P. D. Synergism between Pax-8 and lim-1 in embryonic kidney development. Dev Biol 214, 46–59 (1999).

16. Cirio, M. C. et al. Lhx1 is required for specification of the renal progenitor cell field. PLoS One 6, e18858 (2011).

17. Espiritu, E. B. et al. The Lhx1-Ldb1 complex interacts with Furry to regulate microRNA expression during pronephric kidney development. Sci. Rep. 8, 16029 (2018).

18. Hukriede, N. A. et al. Conserved requirement of Lim1 function for cell movements during gastrulation. Dev Cell 4, 83–94 (2003).

19. Kodjabachian, L. et al. A study of Xlim1 function in the Spemann-Mangold organizer. Int J Dev Biol 45, 209–218 (2001).

20. Shawlot, W. & Behringer, R. R. Requirement for Lim1 in head-organizer function. Nature 374, 425–430 (1995).

21. Chen, L. et al. Ssdp proteins interact with the LIM-domain-binding protein Ldb1 to regulate development. Proc Natl Acad Sci U S A 99, 14320–14325 (2002).

22. van Meyel, D. J., Thomas, J. B. & Agulnick, A. D. Ssdp proteins bind to LIM-interacting co-factors and regulate the activity of LIM-homeodomain protein complexes in vivo. Development 130, 1915–1925 (2003).

23. Bayarsaihan, D., Soto, R. J. & Lukens, L. N. Cloning and characterization of a novel sequence-specific single-stranded-DNA-binding protein. Biochem. J. 331 ( Pt 2, 447–452 (1998).

24. Wang, H., Wang, Z., Tang, Q., Yan, X.-X. & Xu, W. Crystal structure of the LUFS domain of human single-stranded DNA binding Protein 2 (SSBP2). Protein Sci. 28, 788–793 (2019).

25. Wang, H. et al. Crystal structure of human LDB1 in complex with SSBP2. Proc. Natl. Acad. Sci. U. S. A. 117, 1042–1048 (2020).

26. Xu, Z. et al. Single-stranded DNA-binding proteins regulate the abundance of LIM domain and LIM domain-binding proteins. Genes Dev. 21, 942–955 (2007).

27. Wang, Y. et al. SSBP2 is an in vivo tumor suppressor and regulator of LDB1 stability. Oncogene 29, 3044–3053 (2010).

28. Baine, M. J. et al. Transcriptional profiling of peripheral blood mononuclear cells in pancreatic cancer patients identifies novel genes with potential diagnostic utility. PLoS One 6, e17014 (2011).

29. Liu, J.-W. et al. ssDNA-binding protein 2 is frequently hypermethylated and suppresses cell growth in human prostate cancer. Clin. cancer Res. an Off. J. Am. Assoc. Cancer Res. 14, 3754–3760 (2008).

30. Xiao, Y. et al. SSBP2 variants are associated with survival in glioblastoma patients. Clin. cancer Res. an Off. J. Am. Assoc. Cancer Res. 18, 3154–3162 (2012).

31. Kim, H. et al. Single-stranded DNA binding protein 2 expression is associated with patient survival in hepatocellular carcinoma. BMC Cancer 18, 1244 (2018).

32. Moriya, N., Uchiyama, H. & Asashima, M. Induction of Pronephric Tubules by Activin and Retinoic Acid in Presumptive Ectoderm of Xenopus laevis. Dev. Growth Differ. 35, 123–128 (1993).

33. Drews, C., Senkel, S. & Ryffel, G. U. The nephrogenic potential of the transcription factors osr1, osr2, hnf1b, lhx1 and pax8 assessed in Xenopus animal caps. BMC Dev. Biol. 11, 5 (2011).

34. Uochi, T. & Asashima, M. Sequential gene expression during pronephric tubule formation in vitro in Xenopus ectoderm. Dev. Growth Differ. 38, 625–634 (1996).

35. Bowes, J. B. et al. Xenbase: gene expression and improved integration. Nucleic Acids Res. 38, D607–12 (2010).

36. Costello, I. et al. Lhx1 functions together with Otx2, Foxa2, and Ldb1 to govern anterior mesendoderm, node, and midline development. Genes Dev 29, 2108–2122 (2015).

37. Nishioka, N. et al. Ssdp1 regulates head morphogenesis of mouse embryos by activating the Lim1-Ldb1 complex. Development 132, 2535–2546 (2005).

38. Enkhmandakh, B., Makeyev, A. V & Bayarsaihan, D. The role of the proline-rich domain of Ssdp1 in the modular architecture of the vertebrate head organizer. Proc. Natl. Acad. Sci. U. S. A. 103, 11631–11636 (2006).

39. Mukhopadhyay, M. et al. Functional ablation of the mouse Ldb1 gene results in severe patterning defects during gastrulation. Development 130, 495–505 (2003).

40. Huang, S., Johnson, K. E. & Wang, H. Z. Blastomeres show differential fate changes in 8-cell Xenopus laevis embryos that are rotated 90 degrees before first cleavage. Dev. Growth Differ. 40, 189–198 (1998).

41. Dale, L. & Slack, J. M. Fate map for the 32-cell stage of Xenopus laevis. Development 99, 527–551 (1987).

42. Buisson, I., Le Bouffant, R., Futel, M., Riou, J.-F. & Umbhauer, M. Pax8 and Pax2 are specifically required at different steps of Xenopus pronephros development. Dev. Biol. 397, 175–190 (2015).

43. Carroll, T. J., Wallingford, J. B. & Vize, P. D. Dynamic patterns of gene expression in the developing pronephros of Xenopus laevis. Dev Genet 24, 199–207 (1999).

44. Tena, J. J. et al. Odd-skipped genes encode repressors that control kidney development. Dev. Biol. 301, 518–531 (2007).

45. Brennan, H. C., Nijjar, S. & Jones, E. A. The specification and growth factor inducibility of the pronephric glomus in Xenopus laevis. Development 126, 5847– 5856 (1999).

46. Mauch, T. J., Yang, G., Wright, M., Smith, D. & Schoenwolf, G. C. Signals from trunk paraxial mesoderm induce pronephros formation in chick intermediate mesoderm. Dev. Biol. 220, 62–75 (2000).

47. James, R. G. & Schultheiss, T. M. Patterning of the avian intermediate mesoderm by lateral plate and axial tissues. Dev. Biol. 253, 109–124 (2003).

48. Guillaume, R., Bressan, M. & Herzlinger, D. Paraxial mesoderm contributes stromal cells to the developing kidney. Dev. Biol. 329, 169–175 (2009).

49. Cartry, J. et al. Retinoic acid signalling is required for specification of pronephric cell fate. Dev. Biol. 299, 35–51 (2006).

50. Vize, P. D., Jones, E. A. & Pfister, R. Development of the Xenopus pronephric system. Dev. Biol. 171, 531–540 (1995).

51. Smith, J. C. & Watt, F. M. Biochemical specificity of Xenopus notochord. Differentiation 29, 109–115 (1985).

52. Wallingford, J. B., Carroll, T. J. & Vize, P. D. Precocious expression of the Wilms’ tumor gene xWT1 inhibits embryonic kidney development in Xenopus laevis. Dev. Biol. 202, 103–112 (1998).

53. Yasuoka, Y. & Taira, M. LIM homeodomain proteins and associated partners: Then and now. Curr. Top. Dev. Biol. 145, 113–166 (2021).

54. Chan, T. C., Ariizumi, T. & Asashima, M. A model system for organ engineering: transplantation of in vitro induced embryonic kidney. Naturwissenschaften 86, 224–227 (1999).

55. Osafune, K., Nishinakamura, R., Komazaki, S. & Asashima, M. In vitro induction of the pronephric duct in Xenopus explants. Dev. Growth Differ. 44, 161–167 (2002).

56. Hiratani, I., Yamamoto, N., Mochizuki, T., Ohmori, S. & Taira, M. Selective degradation of excess Ldb1 by Rnf12/RLIM confers proper Ldb1 expression levels and Xlim-1/Ldb1 stoichiometry in Xenopus organizer functions. Development 130, 4161–4175 (2003).

57. Lee, B., Lee, S., Agulnick, A. D., Lee, J. W. & Lee, S.-K. Single-stranded DNA binding proteins are required for LIM complexes to induce transcriptionally active chromatin and specify spinal neuronal identities. Development 143, 1721–1731 (2016).

58. Lindenmeyer, M. T. et al. Systematic analysis of a novel human renal glomerulus-enriched gene expression dataset. PLoS One 5, e11545 (2010).

59. Nakano, T., Murata, T., Matsuo, I. & Aizawa, S. OTX2 directly interacts with LIM1 and HNF-3beta. Biochem. Biophys. Res. Commun. 267, 64–70 (2000).

60. Yasuoka, Y. et al. Occupancy of tissue-specific cis-regulatory modules by Otx2 and TLE/Groucho for embryonic head specification. Nat Commun 5, 4322 (2014).

61. Franks, R. G., Wang, C., Levin, J. Z. & Liu, Z. SEUSS, a member of a novel family of plant regulatory proteins, represses floral homeotic gene expression with LEUNIG. Development 129, 253–263 (2002).

62. Nieuwkoop, P. D. & Faber, J. Normal table of Xenopus Laevis. (Garland Publishing, 1994).

63. Gawantka, V. et al. Gene expression screening in Xenopus identifies molecular pathways, predicts gene function and provides a global view of embryonic patterning. Mech. Dev. 77, 95–141 (1998).

64. Tran, U., Pickney, L. M., Ozpolat, B. D. & Wessely, O. Xenopus Bicaudal-C is required for the differentiation of the amphibian pronephros. Dev. Biol. 307, 152– 164 (2007).

65. Miller, R. K. et al. Pronephric tubulogenesis requires Daam1-mediated planar cell polarity signaling. J. Am. Soc. Nephrol. 22, 1654–1664 (2011).

66. Cervino, A. S. et al. Furry is required for cell movements during gastrulation and functionally interacts with NDR1. Sci Rep 11, 6607 (2021).

67. Cizelsky, W., Tata, A., Kühl, M. & Kühl, S. J. The Wnt/JNK signaling target gene alcam is required for embryonic kidney development. Development 141, 2064–2074 (2014).

