## Supplemental Figures and legends for "*Xenopus* Ssbp2 is required for embryonic pronephros morphogenesis and terminal differentiation"

### Supplementary figure legends

**Supplementary Figure S1. Identification of Ldb1-Lhx1 interacting proteins by TAP in kidney-induced animal cap explants.** (**a**) Schematic of the followed procedure. 2-cell stage *Xenopus* embryos were injected with *TAP-LLCA* mRNA, cultured until blastula stage (S8) when animal caps were dissected and cultured for 6 hours in the presence of activin and retinoic acid (RA). Interacting proteins were isolated by TAP of TAP-LLCA and identified by nanoLC/MS/MS. (**b**) List of selected proteins identified in the TAP-LLCA injected sample and absent in the uninjected sample. Highlighted (grey) are proteins expressed in the pronephric kidney (Xenbase) and Ssbp2 (pink) described in this manuscript. In bold are Lhx1 and Ldb1, both part of LLCA. The number of spectral counts (SpC) is indicated for each protein. Full experimental protein list in Supplementary File 1.

**Supplementary Figure S2. Validation of the *ssbp2*-MO specificity.** (**a**) Scheme of the *ssbp2-eGFP* mRNA reporter showing the *ssbp2*-MO target site. (**b-e**) 2-cell stage *Xenopus* embryos were injected into both blastomeres with 1 ng of *ssbp2-eGFP* mRNA in the presence or absence of the standard control morpholino (St-MO) or the *ssbp2*-MO (15 ng). Green fluorescence was analyzed in early gastrula stage embryos (S10.5) to allow *ssbp2-eGFP* mRNA translation. Vegetal views. (**b**) Uninjected embryos (0/40 with fluorescence). (**c**) *ssbp2-eGFP* injected embryos (39/40 with fluorescence). (**d**) *ssbp2-eGFP* + St-MO co-injected embryos (43/43 with fluorescence). (**e**) *ssbp2-eGFP* + *ssbp2*-MO co-injected embryos (2/52 with fluorescence). Note *ssbp2*-MO specifically reduced *ssbp2*-eGFP translation. Two independent experiments were performed. Representative embryos are shown.

**Supplementary Figure S3. Ssbp2 is not essential for establishment of the pronephric field.** 8-cell stage *Xenopus* embryos were injected into a single V2 blastomere as indicated. The uninjected contralateral side was used as internal control. (**a-c**) WISH for *pax8* in late neurula stage embryos (S20). Dorsal views, anterior up. (**a**) Uninjected embryo (3% affected; n = 107; N = 6). (**b**) St-MO 15 ng injected embryo (11% affected; n = 36; N = 2). (**c**) *ssbp2*-MO 15 ng injected embryo (18% affected; n = 89; N = 5). (**d-e**) WISH for *ors2* in mid-neurula stage embryos (S17). Transverse hemi sections, dorsal up. (**d**) Uninjected embryo (7% affected; n = 29; N = 2). (**e**) St-MO 15 ng injected embryo (7% affected; n = 43; N = 3). (**f**) *ssbp2*-MO 15 ng injected embryo (12% affected; n = 60; N = 3). Statistical significance was evaluated using *Chi-square* test. No significant differences were found between groups. N: number of independent experiments, n: number of embryos. Representative embryos are shown.

**Supplementary Figure S4. Ssbp2 depletion affects proximal and distal tubule development.** 8-cell stage *Xenopus* embryos were injected into a single V2 blastomere as indicated. The uninjected contralateral side was used as an internal control. (**a-e**) WISH for *pax8* in 32 stage embryos. (**a**) Uninjected embryo

(5% affected, n = 40, N = 2). (**b,c**) St-MO 15 ng injected embryo (10% affected, n = 41, N = 2). (**d,e**) *ssbp2*-MO 15 ng injected embryo (44% affected, n = 54, N = 3). (**f-j**) WISH for *hoxb7* in stage 32 embryos. (**f**) Uninjected embryo (10% affected, n = 40, N = 2). (**g,h**) St-MO 15 ng injected embryo (31% affected, n = 35, N = 2). (**i,j**) *ssbp2*-MO 15 ng injected embryo (68% affected, n = 38, N = 2). Magnifications of the pronephric tubules enclosed by the black squares are shown in the left-bottom corner. Statistical significance was evaluated using *Chi-square* test (****p < 0.0001; ***p < 0.001). Representative embryos are shown. (**k**) Quantification of the tubule length in the most anterior *hoxb7* expression domain revealed by WISH (dotted red lines in *f-j*). The ratio between the injected and the control side is shown. Data in graph is presented as mean and standard deviation. Each point represents a single embryo. Statistical significance was evaluated using *Kruskal–Wallis* test and *Dunn's* multiple comparisons test (**** p < 0.0001; ***p < 0.001). * represent the comparison to the uninjected group and † represents the comparison to the St-MO injected group. N: number of independent experiments, n: number of embryos.

### Supplementary Figure S1.

a

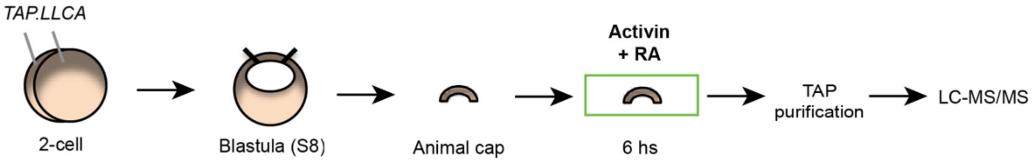


b

| Identified Proteins | Accession Number | TAP.LLCA (SpC) | Uninjected (SpC) |
|---|---|---|---|
| pyruvate carboxylase, gene 2 | AAI35599 | 71 | 0 |
| propionyl CoA carboxylase, alpha polypeptide | NP_001089298 | 57 | 0 |
| methylcrotonoyl-CoA carboxylase 1 (alpha) | NP_001086068 | 33 | 0 |
| **LIM-domain-binding protein 1b** | BAE95405 | 22 | 0 |
| propionyl-CoA carboxylase subunit beta L homeolog | AAH61665 | 21 | 0 |
| tubulin alpha like 3, gene 1 L homeolog | AAH84938 | 17 | 0 |
| methylcrotonyl-CoA carboxylase subunit 2 S homeolog | AAI29636 | 17 | 0 |
| Keratin, type I cytoskeletal 18-A | P08802 | 12 | 0 |
| polyubiquitin-B precursor | AAI26016 | 12 | 0 |
| heat shock 70kDa protein 1-like | NP_001080068 | 9 | 0 |
| tubulin beta-4 chain | NP_001080566 | 9 | 0 |
| receptor for activated C kinase 1 S homeolog | NP_001086465 | 7 | 0 |
| unnamed protein product | CBF67870 | 7 | 0 |
| NADH:ubiquinone oxidoreductase subunit A12 | AAH84520 | 7 | 0 |
| heat shock protein family D (Hsp60) member 1 | AAI3584 | 6 | 0 |
| heat shock protein 90kDa alpha, class B member 1 | NP_001086624 | 5 | 0 |
| ribosomal protein L23 | NP_001085921 | 4 | 0 |
| ATP synthase, H+ transporting, mitochondrial F1 complex, gamma polypeptide 1 | NP_001080481 | 4 | 0 |
| MGC82602 protein | NP_001085608 | 4 | 0 |
| Vitellogenin-B2 | P19011 | 4 | 0 |
| solute carrier family 25, member 11 L homeolog | AAI23334 | 4 | 0 |
| arginase 1 | NP_001080417 | 3 | 0 |
| single-stranded DNA binding protein 2 | NP_001080347 | 3 | 0 |
| electron transfer flavoprotein subunit alpha S homeolog | NP_001090035 | 3 | 0 |
| NADH dehydrogenase (ubiquinone) 1 alpha subcomplex, 6 | NP_001088970 | 3 | 0 |
| NADH dehydrogenase (ubiquinone) Fe-S protein 4 | NP_001087349 | 3 | 0 |
| **LIM homeobox 1 protein** | AAI35732 | 3 | 0 |
| 40S ribosomal protein S7 | P02362 | 3 | 0 |
| mitochondrial ATP synthase beta subunit | NP_001080126 | 2 | 0 |
| ribosomal protein S14 | AAI35233 | 2 | 0 |
| ATP synthase, H+ transporting, mitochondrial F1F0 complex, subunit e | AAI67416 | 2 | 0 |
| tubb4 protein | AAH64270 | 2 | 0 |
| fish-egg lectin | AAI53784 | 2 | 0 |
| chaperonin containing TCP1, subunit 6A (zeta 1) | NP_001086080 | 2 | 0 |
| peroxiredoxin 1 L homeolog | NP_001085178 | 2 | 0 |
| phosphofructokinase, muscle | NP_001086921 | 2 | 0 |
| proteasome alpha 6 subunit | NP_001086785 | 2 | 0 |
| ribosomal protein S18 | NP_001084747 | 2 | 0 |
| ribosomal protein L30 | NP_001080621 | 2 | 0 |
| hyaluronan binding protein 4 L homeolog | NP_001083218 | 2 | 0 |
| glyceraldehyde 3-phosphate dehydrogenase [Xenopus laevis] | AAA84422 | 2 | 0 |
| SPT2, Suppressor of Ty, domain containing 1 [Xenopus (Silurana) tropicalis] | AAI21681 | 2 | 0 |

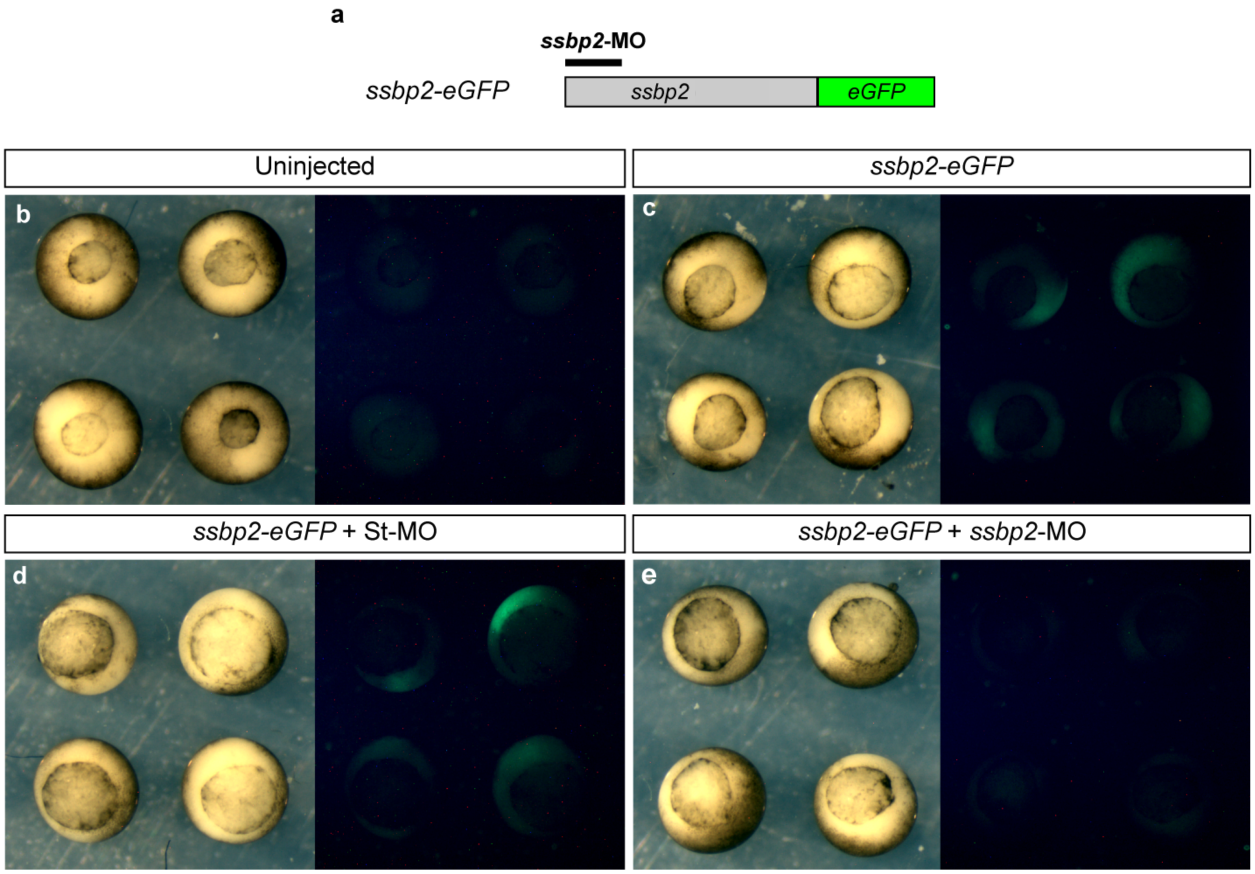


**Supplementary Figure S2.**

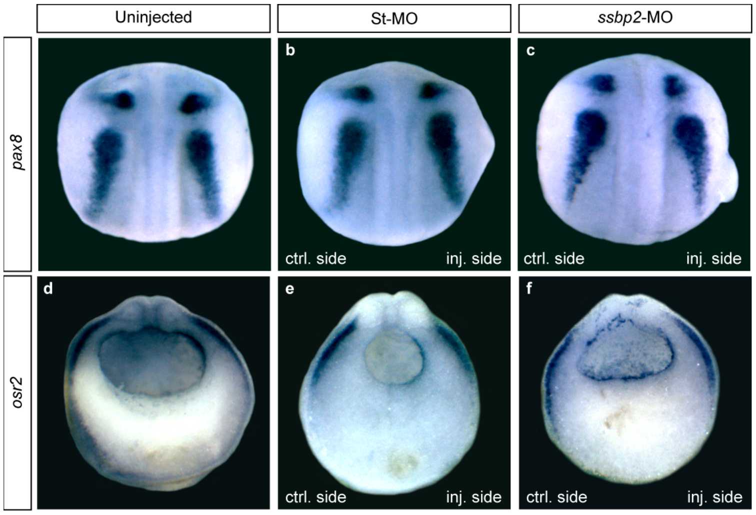


**Supplementary Figure S3.**

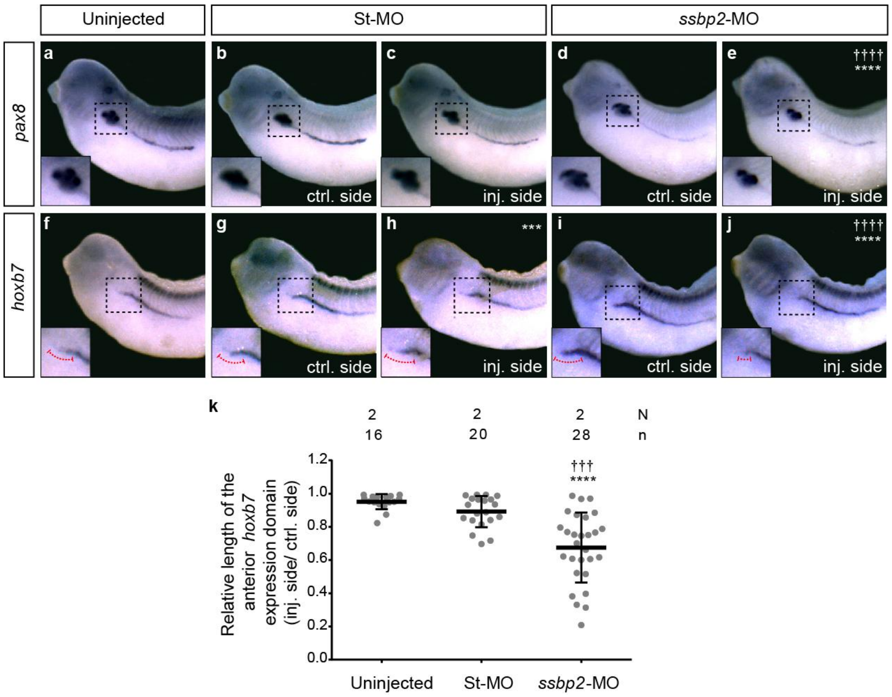


**Supplementary Figure S4.**
